# Low dimensionality of phenotypic space as an emergent property of coordinated teams in biological regulatory networks

**DOI:** 10.1101/2023.02.03.526930

**Authors:** Kishore Hari, Pradyumna Harlapur, Aashna Saxena, Kushal Haldar, Aishwarya Girish, Tanisha Malpani, Herbert Levine, Mohit Kumar Jolly

## Abstract

Biological networks driving cell-fate decisions involve complex interactions, but they often give rise to only a few phenotypes, thus exhibiting low-dimensional dynamics. The network design principles that govern such cell-fate canalization remain unclear. Here, we investigate networks across diverse biological contexts– Epithelial-Mesenchymal Transition, Small Cell Lung Cancer, and Gonadal cell-fate determination – to reveal that the presence of two mutually antagonistic, well-coordinated teams of nodes leads to low-dimensional phenotypic space such that the first principal component (PC1) axis can capture most of the variance. Further analysis of artificial team-based networks and random counterparts of biological networks reveals that the principal component decomposition is determined by the team strength within these networks, demonstrating how the underlying network structure governs PC1 variance. The presence of low dimensionality in corresponding transcriptomic data confirms the applicability of our observations. We propose that team-based topology in biological networks are critical for generating a cell-fate canalization landscape.

## 1 Introduction

Cellular decision-making is driven by the complex non-linear dynamics of underlying gene regulatory networks. Despite involving a large number of interacting components, these networks often enable only a limited number of eventual phenotypes/cell types [1, 2]. For example, in differentiation events, the cells move from a high-entropy progenitor phenotype to a low-entropy differentiated phenotype [3]. Such changes in entropy suggest a large-scale modification of gene expression patterns during differentiation. Another well-studied example of cellular decision-making is epithelial-mesenchymal plasticity (EMP), a crucial cellular capability implicated in development, regeneration, and cancer metastasis [4]. Populations capable of undergoing EMP show two major phenotypes: Epithelial (E) and Mesenchymal (M). The E and M phenotypes have mutually exclusive biochemical signatures, and their master regulators inhibit each other [5, 6, 7], reminiscent of the presence of a “toggle switch” between master regulators at many developmental cell-fate bifurcation events [8]. A minor fraction of populations also exhibit hybrid E/M phenotypes with mixed biochemical signatures to varying extents [9, 10, 11, 12]. The hybrid E/M phenotypes have complex non-uniform biochemical compositions compared to the E and M phenotypes, making them hard to identify and therapeutically target [13]. Both progenitor phenotypes and the hybrid E/M phenotypes lack the coherent expression profiles of differentiated cell types and the E and M phenotypes, respectively. In other words, these phenotypes exist in a higher dimensional space than the E and M states, even as the E and M states can be reduced to the composite axis composed of canonical Epithelial and Mesenchymal genes [14, 15]).

A recent analysis of EMP and other related regulatory networks recently suggested that most of the variance in phenotypic space could be explained by the first principal component (PC1) axis [16], suggesting that many cell-fate decision systems, including those of EMP, operate in a low-dimensional space. However, what underlying network features allow for a high PC1 variance has been unclear. In our previous work [5], we demonstrated a seemingly different phenomenon in diverse networks of EMP. Despite their complexity, their repertoire of emergent steady states is limited; thus, most of the phenotypic space converges to Epithelial or Mesenchymal states. We demonstrated that this convergence was a direct consequence of underlying network topology. These networks comprised well-coordinated “teams” of nodes, such that the nodes belonging to the same team effectively activate each other, while those across teams inhibited one another. How connected (or not) are the concepts of “teams” and PC1 variance remains unclear.

Here, we reconcile the observations related to PC1 variance and the presence of “teams” of nodes as a design principle underlying robust phenotypic landscapes. Investigating regulatory networks across biological contexts – EMP, small cell lung cancer, and gonadal cell-fate determination –we show that networks forming strong teams show a high PC1 dominance, a property that is lost upon the weakening of teams by deletion of specific edges and/or swapping edges within the network. The weaker the team strength, the more the number of principal components needed to explain the underlying variance, and thus, the higher the dimensionality of phenotypic space. To generalize our results further, we generated artificial team-based networks of varying densities to highlight how underlying network topology governs the team strength and, consequently, the PC1 variance. Finally, we demonstrate this low dimensionality trait in experimental RNA-sequencing data, a behavior specific to relevant genes forming “teams” but not for housekeeping genes. Overall, our results elucidate how the presence of well-coordinated “teams” of nodes in underlying networks can lead to low dimensionality of phenotypic space and concomitant cell-fate canalization during decision-making.

## 2 Results

### 2.1 Teams of nodes in EMP network and the dimensionality of the steady state space

We start by analyzing a 22 node 82 edge (22N 82E) EMP network [17] (Figure 1A, i). Each node in this network is either a transcription factor or a micro-RNA. Each edge represents the regulation (either transcriptional or post-transcriptional) of the target node (gene) by the source node. These regulations can be activating or inhibiting. As demonstrated in our previous work, the influence matrix generated from this network (which takes a weighted sum of direct and indirect interactions between each node pair) can be effectively considered a higher-order toggle switch (Figure 1A, ii). We refer to each “supernode” (collection of similarly behaving nodes) in this toggle switch as a “team” consisting of a set of nodes positively influencing each other and negatively influencing the nodes of the other team (Figure 1A, ii).

**Figure 1:**
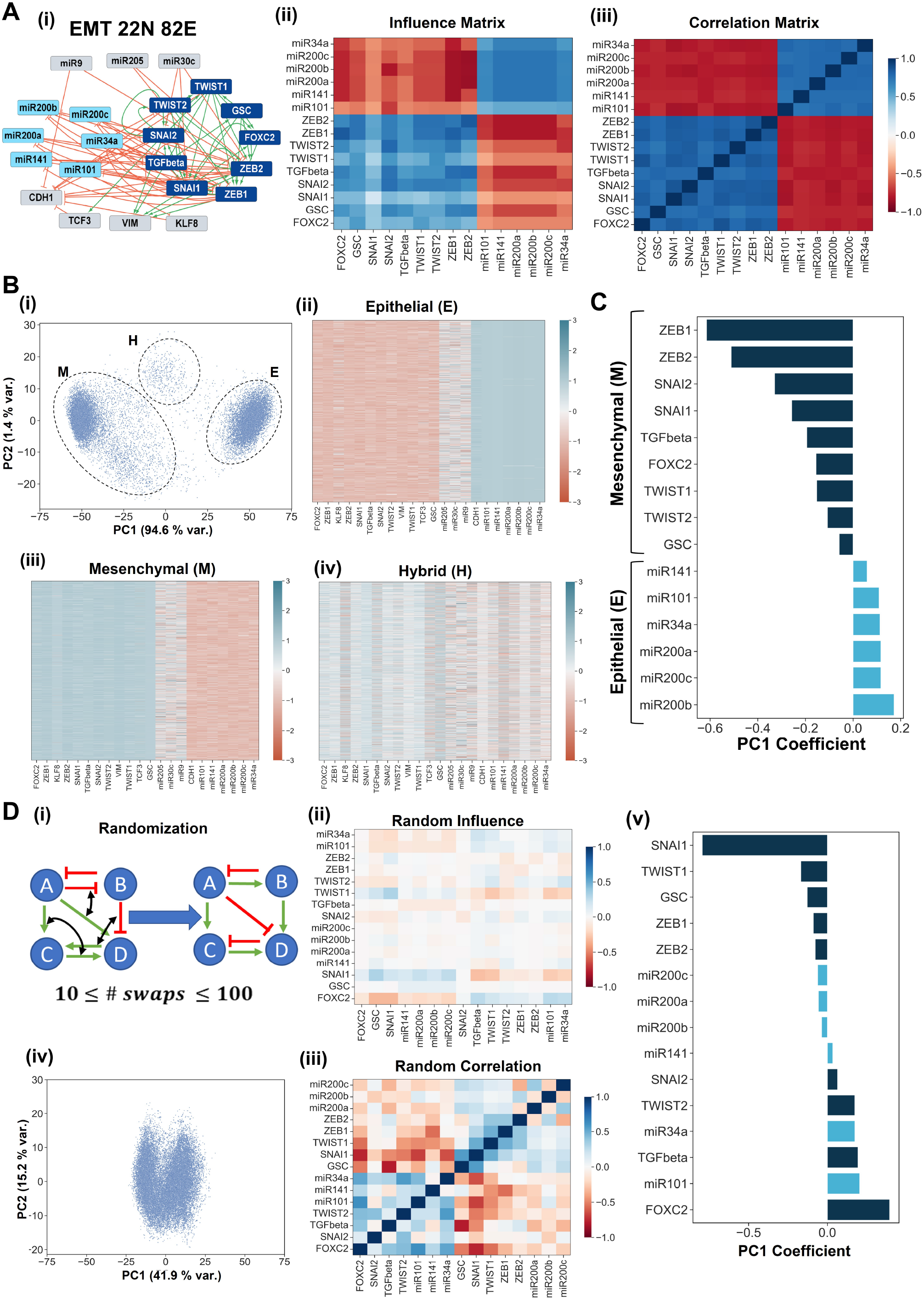
Structural similarities between teams and PC1. **A** The network diagram (i) and the influence matrix (ii) for the 22N 82E EMP network. (iii) Correlation matrix depicting the pairwise correlations between the node expression levels across all parameter sets in RACIPE. **B** (i) Scatterplot mapping the solutions generated from RACIPE on the axes of PC1 and PC2. (ii-iv) Heatmaps depicting the expression levels of the nodes of the network for each individual cluster seen in (i). **C** The loadings of the nodes for each PC1 axis. The colors of the bars represent the team identity of the nodes. **D** (i) Depiction of network randomization. (ii) Influence matrix of a random network. (iii) Correlation matrix generated for a random network. (iv) Scatterplot similar to C (i) but for a random network. (v) Similar to D but for a random network.

We simulate this network over many kinetic parameter sets using the software tool RACIPE [17]. Briefly, RACIPE first generates a system of coupled Ordinary Differential Equations (ODEs) for a particular topology of GRN. It then randomly samples values for each parameter of the ODE system from a corresponding self-consistently determined physiological range to generate an ensemble of parameter sets. RACIPE then simulates the ODEs for each parameter set over multiple initial conditions to identify the steady states. Despite the heterogeneity in the parameter sets, the expression levels of the nodes of the network were well correlated with each other. Notably, the team structure seen in the influence matrix (obtained directly from the network structure, without any simulations) was reflected in the pairwise correlation matrix obtained from dynamical simulations, where the epithelial nodes (such as miR200c, miR200b) are strongly positively correlated with the epithelial nodes and negatively with the mesenchymal nodes (such as ZEB1, SNAIL) (Figure 1A, iii). We then performed a principal component analysis (PCA) over the collection of all steady states obtained for this network. When the solutions are plotted along the first and second principal component axes (PC1 and PC2, respectively), we found the steady states separated into 3 clusters (Figure 1B, i). Analyzing the expression patterns of the steady states forming these clusters revealed that the two prominent clusters were the epithelial and mesenchymal phenotypes. In contrast, the smaller cluster showed a mixed expression of epithelial (E) and mesenchymal (M) nodes and hence could be classified as a hybrid (E/M) cluster (Figure 1B ii-iv). Furthermore, we observed that the PC1 was enough to distinguish between the Epithelia and Mesenchymal steady states (Figure 1B, i). Given our previous understanding of the connection between teams of nodes and the emergent phenotypes, we hypothesized that the two-team structure in the network could drive the relative weights of node expressions that make up the PC1 axis. To verify this hypothesis, we looked at the composition of the PC1 axis (using the node coefficients of PC1) and compared it against the composition of teams (Figure 1C, nodes belonging to different teams are colored differently). In line with our hypothesis, we see that the PC1 contributions from nodes belonging to different teams have opposite signs and that those belonging to the same team have the same sign. Together, these results support the hypothesis that team structure leads to PC1 dominance, wherein, it explains most of the variance in steady-state space.

An immediate implication of the above hypothesis would be that in the absence of strong teams, the PC1 axis would no longer be able to explain the majority of the variance in the steady-state space. To test the validity of this implication, we generated random networks by swapping many pairs of edges in the network (Figure 1D, i)). We have previously shown that such randomization can disrupt the team structure and thus the stability and frequency of E, M, and H phenotypes (Figure 1D, ii). We simulated each generated random network using RACIPE and performed PCA on the steady-state values given by the simulations. Unlike what happens in the wild-type EMP network, in the random networks the PC1 and PC2 axes could not separate the steady states into distinct clusters (Figure 1D, iii, iv). We also found that the classification of the nodes based on their PC1 coefficients did not match their teams’ classification (Figure 1E, v). Therefore, the presence of two mutually inhibiting teams in WT networks appears to be necessary to enable low dimensionality in the emergent phenotypic space.

### 2.2 Team strength correlates with PC1 variance and dimensionality of random networks

Next, we compared the percentage variance explained along the PC1 axis for WT networks against that of random networks. We found that this measure was much lower in random networks than in the WT networks. In the wild-type network, PC1 alone explained nearly 90% of the variance (Figure 2A, i). A scatter plot between the team strength of the random and WT networks against the variance explained by PC1 (Figure 2A, ii) revealed a strongly positive correlation between the two measures, directly suggesting that an increase in team strength can lead to a corresponding increase in the variance explained by PC1. An important observation here is the sigmoidal nature of the data. After a team strength of about 0.25-3, the variance explained by PC1 starts to saturate.

**Figure 2:**
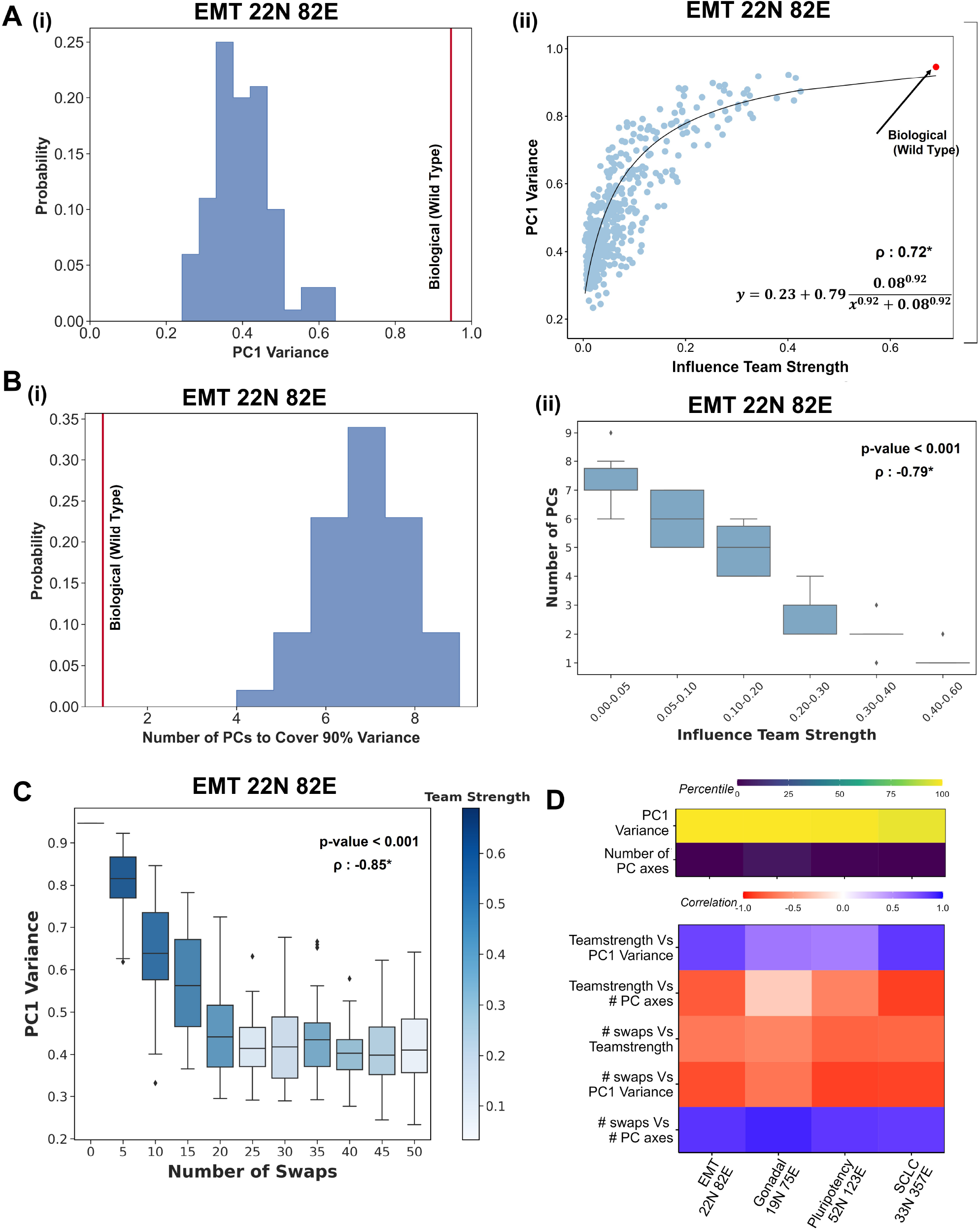
Strong teams lead to a low-dimensional steady-state space.. **A** (i) Histogram depicting the variance explained by PC1 for random networks. EMP network is represented with a vertical red line. (ii) Scatterplot depicting the dependence of variance explained by PC1 on the team strength. Each point is a random network. The biological network has been pointed out. The Spearman correlation coefficient, *ρ*, is reported, with * indicating a *p − value <* 0.05. **B** (i) Histogram depicting the number of PCs required to explain 90% variance in random networks. EMP network is represented with a vertical red line. (ii) Boxplot depicting the dependence of the number of PCs required to explain 90% of the variance in steady states on the team strength. The p-value for one-way ANOVA and the Spearman correlation coefficient (*ρ*) have been reported, with * indicating a *p − value <* 0.05. **C** PC1 variance as a function of the number of swaps used to generate random networks. **D** Heatmap depicting the percentile of Biological networks for PC1 variance, number of PC axes in random networks, and correlation strength between different metrics analyzed here. The p-value for one-way ANOVA and the Spearman correlation coefficient (*ρ*) have been reported, with * indicating a *p − value <* 0.05.

We next wanted to estimate the dimensionality of the steady-state space emergent from these networks. More than 90% of the variance in the phenotypic space of the EMP network is explained by the first principal component axis, making it effectively one-dimensional [16]. We then asked how many principal components are necessary for explaining 90% variance in random networks. While the WT network only requires one principal component (PC1), the random networks require a higher number of PCs, ranging from 7 to 13 out of the 22 axes (Figure 2B, i), demonstrating the high-dimensionality of the phenotypic space of random networks as compared to the biological networks. The number of PCs also showed a strong negative correlation with team strength (Figure 2B, ii), showing that the presence of strong teams reduces the dimensionality of the steady-state space.

To further emphasize the role of the network topology, we compared random networks generated from the WT over a range of edge swaps. Swapping the edges changes the topological features of the biological network. The more the number of swaps, the greater the changes in the topology, which should be reflected in the emergent behavior. Indeed, we find that the PC1 variance decreases with increasing number of swaps (Figure 2C) along with the team strength (color of boxes in Figure 2C).

### 2.3 Strong teams underlie the low-dimensionality of phenotypic space in multiple contexts of cell-fate decisions

Our observations establish that strong teams underly the PC1 dominance in the 22N 82E EMP network. We then asked if the aforementioned results hold in other biological contexts. We also investigated three additional regulatory networks underlying cell-fate decisions: SCLC, pluripotency, and gonadal fate determination (Figure S1A) [18, 19, 20]. The SCLC network has been shown to regulate the phenotypic heterogeneity underlying small-cell lung cancer, allowing the cells to switch between neuroendocrine (NE) and non-neuroendocrine (non-NE) phenotypes. The Pluripotency network describes the differentiation of hESCs. The network was constructed to identify pathways in which iPSCs can be achieved. The gonadal fate determination network describes the cell-fate determination between Sertoli and granulosa cell types, events in the early stages of the development of male and female fates respectively. These three networks display a spectrum of characteristics. The Pluripotency network has hub genes (NANOG, SOX2, OCT4) with high out-degree (i.e., these nodes regulate multiple genes). SCLC and Pluripotency networks have relatively lower team strengths (0.25 and 0.2), which puts them near the border of saturation in the PC1 variance vs team strength plot in Figure 2B. All three networks show a clear dominance over corresponding random networks, as shown by the percentile plots in Figure 2D. The gonadal and SCLC networks show a clear team structure in their influence matrices (Figure 3A, i, ii). Such a team structure is relatively harder to discern in the iPSC influence matrix (Figure 3A, iii). However, our clustering-based method of identifying teams allowed us to classify the nodes into two teams. This classification agreed with the biological function of these molecules and with the groups seen in the correlation matrix (Figure S1B). For all three networks, the nodes of different teams showed opposite signs in PC1 loading (Figure 3B). This finding is particularly noteworthy for the pluripotency network, given its weak team strength. The PC1 could separate the steady states into clusters, defining clear phenotypes for all three networks (Figure S2). These three networks also maintained the correlations among PC1 variance, number of PC axes, team strength, and number of swaps observed in the EMP network (Figure 2D). Furthermore, we found that the sigmoidal relationship between the team strength and PC1 variance seen for the EMP network (Figure 2A, ii) was also maintained for these three networks (Figure 3C). Lastly, we also confirmed that the inverse trend between team strength and the number of PCs needed to explain 90% variance in steady-state space was also seen for these three networks. However, the monotonicity of the relationship between team strength and the number of PCs was broken for the Pluripotency network, perhaps because it has the weakest team composition of the three. Furthermore, as the number of swaps increased, the PC1 variance for the random networks and the team strength of the random networks decreased simultaneously. (Figure 3D). Together, these results assert that strong teams universally regulate the dimensionality of the emergent phenotypic space of cell-fate decision-making networks.

**Figure 3:**
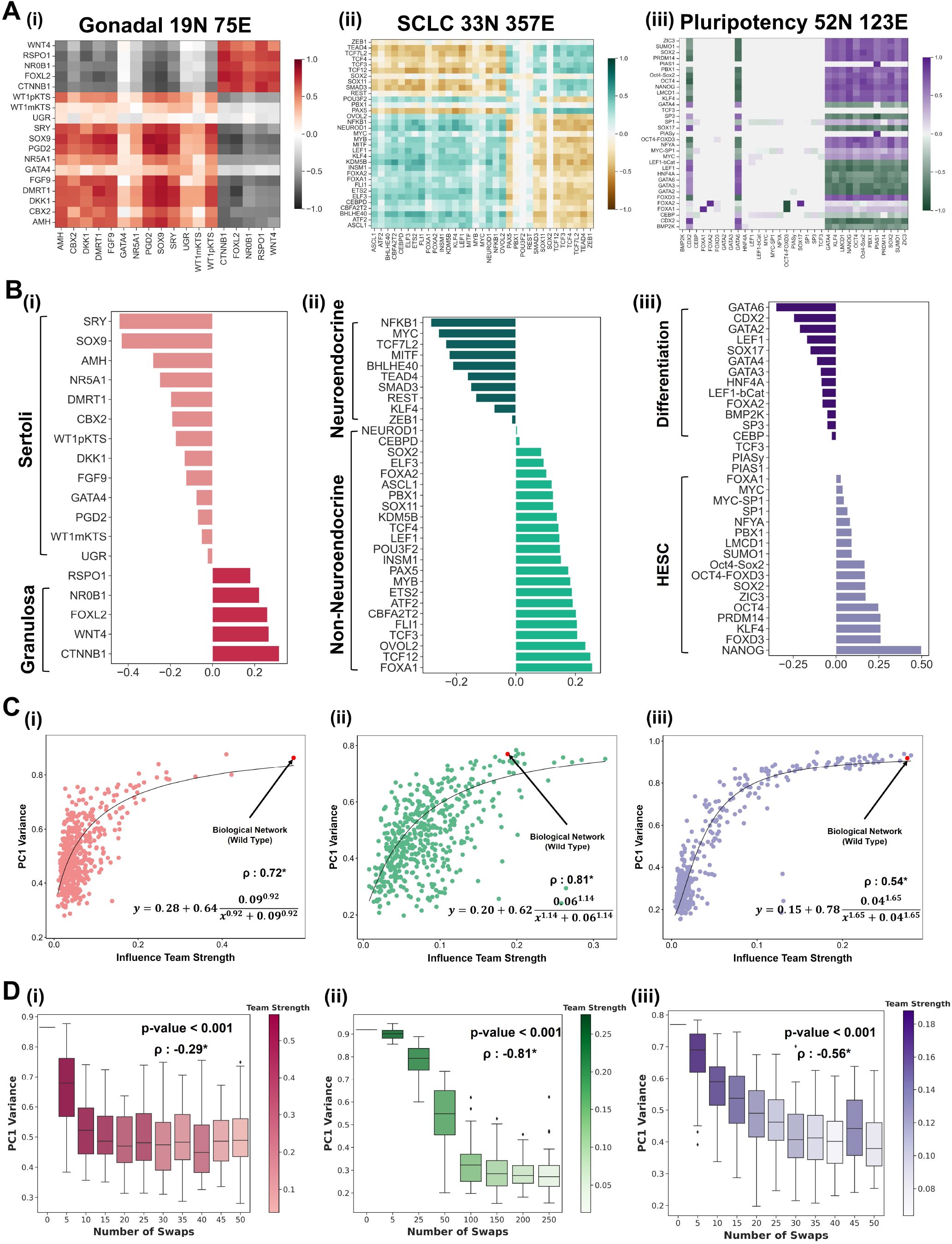
Team structure leads to low-dimensionality of steady-state space in SCLC, iPSC, and gonadal cell-fate decision networks. **A** Influence matrices for (i) Gonadal, (ii) SCLC and (iii) iPSC network. **B** The loadings of the nodes for each PC1 axis for (i) Gonadal, (ii) SCLC, and (iii) iPSC network. The colors of the bars represent the team identity of the nodes. **C** Scatterplot depicting the dependence of variance explained by PC1 on the team strength for (i) Gonadal, (ii) SCLC, and (iii) iPSC network. Each point is a random network. The biological network has been pointed out. The Spearman correlation coefficient (*ρ*) is reported, with * indicating a *p − value <* 0.05. **D** Boxplot depicting the dependence of PC1 Variance on the number of swaps for (i) Gonadal, (ii) SCLC, and (iii) iPSC network. The p-value for one-way ANOVA and the Spearman correlation coefficient (*ρ*) have been reported, with * indicating a *p − value <* 0.05.

### 2.4 Reducing team strength increases steady state-space dimensionality

Our random network analysis suggests that the team structure in biological networks gives rise to low-dimensional phenotypic space due to the strong phenotypic reinforcement provided by the teams. To establish causation, we wanted to directly examine how weakening teams would affect dimensionality. Since teams are formed due to the strong interactions within the network, we chose to disrupt teams by disconnecting nodes from the network. While the random network generation by swapping edges reduces teamstrength by introducing inconsistent links such as inhibitions within teams and activations across teams, this method maintains the consistency of the edges while reducing the density of the edges. Biologically, disconnected nodes can imply mutations or nodes erroneously included in the network.

We increasingly disconnected the existing nodes in the network - one at a time - by removing their incoming and outcoming edges and adding either a self-activation (SA) or self-inhibition (SI) on them. Given the many possible sequences of removing nodes one at a time, we repeated this analysis 100 times for each iteration (i.e., for a fixed number of disconnected nodes). As expected, we correspondingly see a decrease in the teamstrength of the network with an increasing number of nodes disconnected (Figure 4A, i). We find that for the EMT 22 node network, the PC1 variance showed a sigmoidal decline as the number of disconnected nodes increased (Figure 4A, ii). The means of the PC1 variance distributions of these networks showed a sigmoidal decrease with an increasing number of nodes disconnected (Figure 4A, iii). Adding SI to the disconnected nodes leads to a higher threshold value than adding SA. A possible explanation could be the noise suppression characteristics of the self-inhibition motif, in contrast to the noise-amplifying behavior of the self-activation motif. Thus, the self-activation-based curve can represent the maximum resilience offered by the team structure against erroneous inclusion of unrelated genes in genesets corresponding to cell-fate decision-making. Note that the number of nodes at which the PC1 variance starts to decrease corresponds to a mean team strength of around 0.3. A similar trend could also be seen for the gonadal network: a decrease in the team strength (Figure 4B, i), a sigmoidal decrease in the PC1 variance with an increase in the number of nodes disconnected (Figure 4B ii-iii). Note that the PC1 variance remains unchanged for a larger percentage of node disconnects in Gonadal network (6*/*19 = 32%) compared to the EMP network (4*/*22 = 19%). This contrast can be explained by the reduction rate of team strengths in both cases. EMP network sees a much sharper decrease in teamstrength against the number of nodes disconnected (Figure 4A iii) compared to the Gonadal network (Figure 4B iii). Together, these results help establish a causal relationship between the team strength and the low dimensionality of the phenotypic space emergent from simulated data.

**Figure 4:**
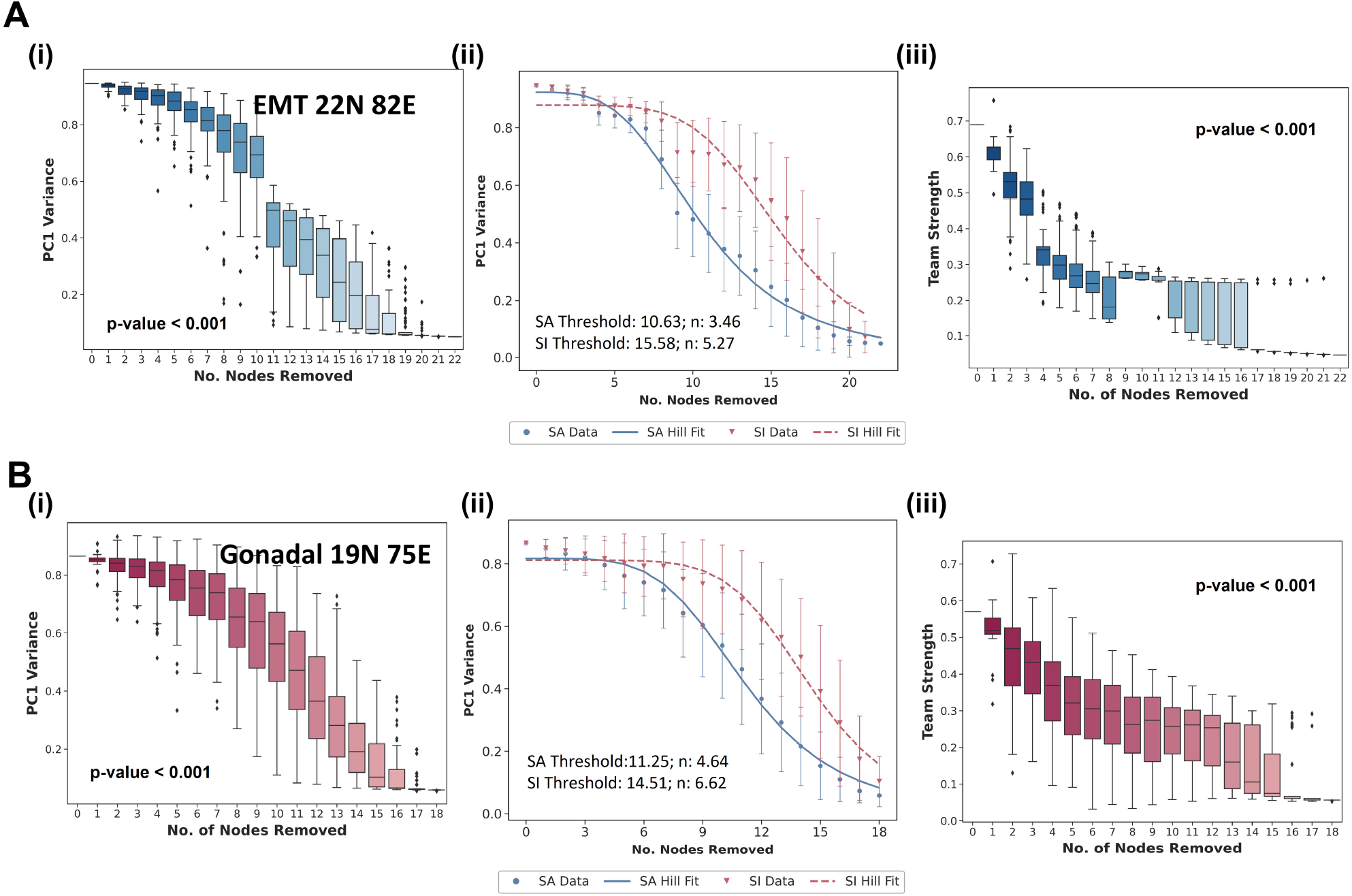
Weakening teams in the Biological networks increases dimensionality in the steady state space. Networks generated by removing *n* (0≤*n*≤*totalnumber of nodes*) nodes in the network and replacing them with *n* disconnected nodes with self-activation. 50 networks generated for each *n*. **A**. For the 22N EMT network, (i) Boxplot describing the change in PC1 Variance with the number of nodes replaced with self-activating disconnected nodes, (ii) Mean ± sd PC1 variance against the number of nodes replaced with self-activating (blue) and self-inhibiting (red) disconnected nodes, (iii) Team strength against the number of nodes replaced. Fits to sigmoidal curve are shown using blue and red lines in (ii) and the corresponding parameters are reported in the plot. **B**. Same as A but for Gonadal cell-fate network.

### 2.5 Low-dimensionality and indicators of team structure in transcript-tomic data

Our analysis so far has proposed team structure in the underlying gene regulatory network as a mechanism behind the low dimensionality of the simulated expression matrices. We then proceeded to test the dimensionality of transcriptomic data for cells governed by either the EMT or SCLC cell-fate networks and asked if the presence of teams can be perceived in these RNA sequencing datasets and if this presence leads to low dimensionality. First, we investigate the cell-line expression data from CCLE [21], for transcriptional signatures of EMT seen in carcinomas as well as the NE/non-NE axis of cellular plasticity observed in SCLC [22, 23, 24]. We analyzed the SCLC gene signature with the SCLC cell lines in the CCLE and the EMT signature with other cell lines.

Similar to the simulation data; the pairwise correlation matrix obtained from gene expression showed a clear separation of two teams in all data sets for the correspondingly appropriate gene lists - SCLC gene sets for the SCLC cell lines (Figure 5A, i) and EMT gene sets for non-SCLC cell lines (Figure 5B, i). Further endorsing our simulation results, the coefficients of the genes in the first principal component axis reflect their team assignment (Figure 5A, ii; 5B, ii). Furthermore, the first principal component explained a significant part of the variance in the expression data (Figure 5A, iii; 5B, iii). The different cell lines clustered by their corresponding NE and EMT scores **Citation**. This feature of low dimensionality is unique to these relevant gene lists and is lost for a random set of genes (Figure S3).

**Figure 5:**
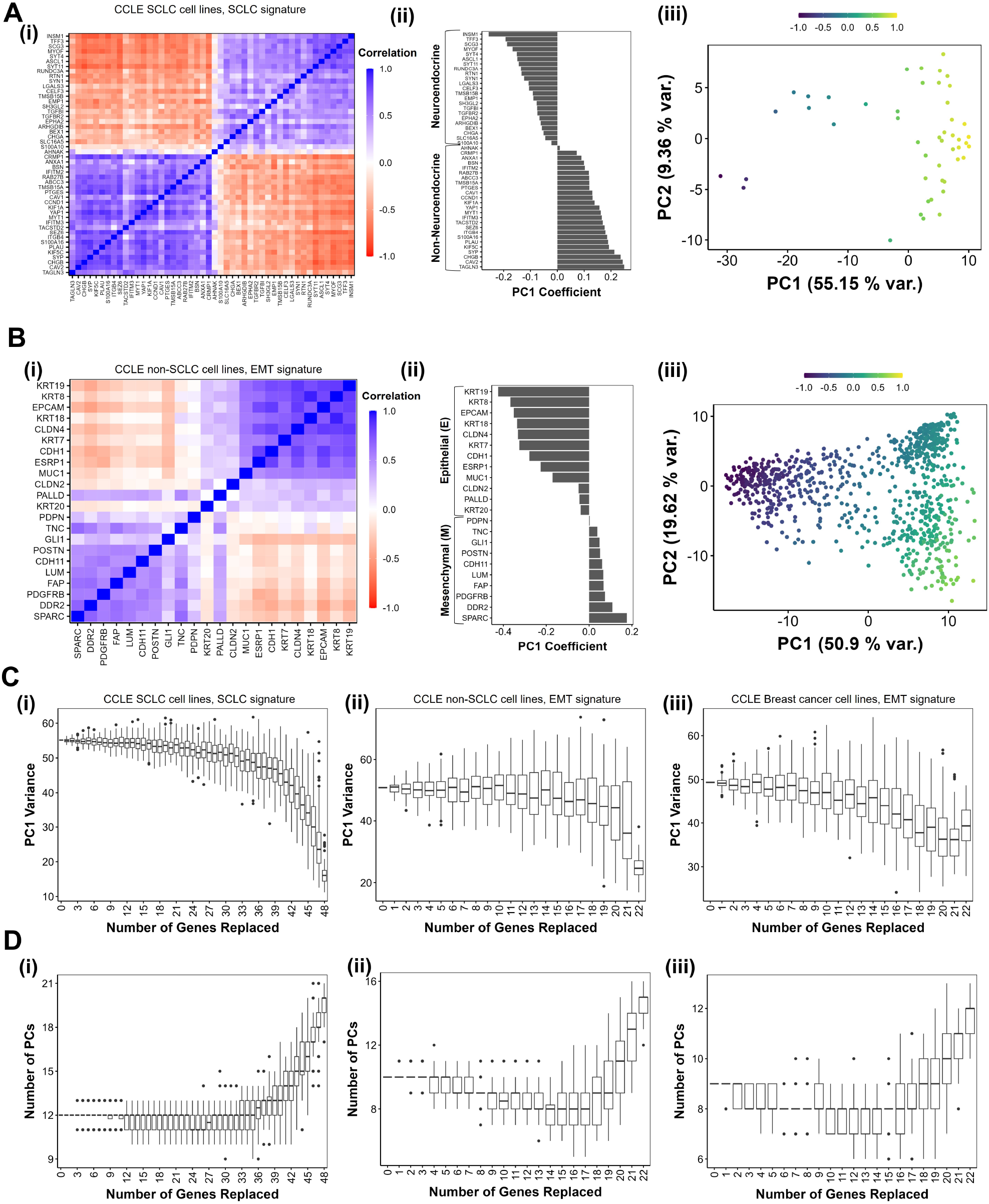
Low-dimensionality in CCLE datasets. **A**. (i) Pairwise correlation of expression vectors of the 32 genes of SCLC network in the CCLE SCLC cell-line data. (ii). Coefficients of first principle component axis for the geneset-cell line combination in (i). (iii) Scatter plot of CCLE expression data on the PC 1 and 2 axis. Each cell line is colored using a phenotypic score calculated as a difference in the cumulative, max-normalized expression of NE and non-NE genes. **B**. Same as A, but for CCLE non-SCLC cell-lines for EMT signature. **C**. PC1 Variance against the number of genes replaced with housekeeping genes for (i) CCLE SCLC cell-lines with SCLC gene signature, (ii) CCLE non-SCLC cell-lines with EMT signature and (iii) CCLE Breast cancer cell lines with EMT signature. **D**. Same as C but for number of PCs needed to explain 90% Variance.

We then asked how the dimensionality of these datasets would change when network components were increasingly replaced by irrelevant genes, thereby forming a somewhat erroneous geneset. For a geneset of length N, we replaced n genes at a time with a set of housekeeping genes [25] and calculated the principal components for the new set of N genes. We repeat this exercise 100 times for each value of n (Figure 5C, D). First, note that PC1 variance decreases with an increasing number of genes replaced with housekeeping genes, while the number of PC axes required to explain 90% variance increases. Both the results suggest that the dimensionality of these datasets increases with increased error in the geneset composition, as expected.

Interestingly, the increase in dimensionality is minimal till nearly 80% of the genes are replaced with housekeeping genes. This behavior is evidence of the strong control structure underlying these genesets. As compared to a similar experiment done on simulated data (Figure 4), at first glance, it would seem like the transcriptomic data shows higher resilience than simulated data. A possible explanation for this difference is that our simulations disrupted the team structure but in transcriptomic data analysis we are simply studying a different set of genes than those necessarily participating in teams. We demonstrate this difference by performing a similar analysis on the simulated expression matrices (Figure S4). Indeed, when the team structure is not disrupted, we see a higher resilience to node replacement than when the team structure is disrupted (Figure S4 and Figure 4). Thus, these results indicate that the low dimensionality of the transcriptomic data corresponding to SCLC/EMP cell-fate decisions robustly witnessed in RNA-seq data.

### 2.6 Artificial networks reveal edge density as a key variable affecting team strength and dimensionality

Our results so far have been obtained for biological networks and randomized networks generated therefrom. While these networks feature mutually inhibiting teams of nodes and low dimensionality of the emergent phenotypic space, topological structures other than teams may have a significant role. Therefore, to further elucidate the causative connection between team structure and low dimensionality, we generated networks of size 10 with pre-defined team structure and with varying edge densities (Figure 6A). By pre-defining the team structure, we ensure that the edges between the nodes belonging to the same team are only activating, while those between nodes belonging to different teams are always inhibiting. Since the influence between a pair of nodes is a weighted sum of the paths running between them, we hypothesized that by controlling the edge density, we could control the number of paths between nodes and, hence, the team strength. Each edge in the network has an equal probability of being filled, and the probability value determines the network density. We used the Boolean formalism [14] to simulate these networks for a reduced computational cost to be able to probe a higher resolution of density, teamstrength, and the number of swaps.

**Figure 6:**
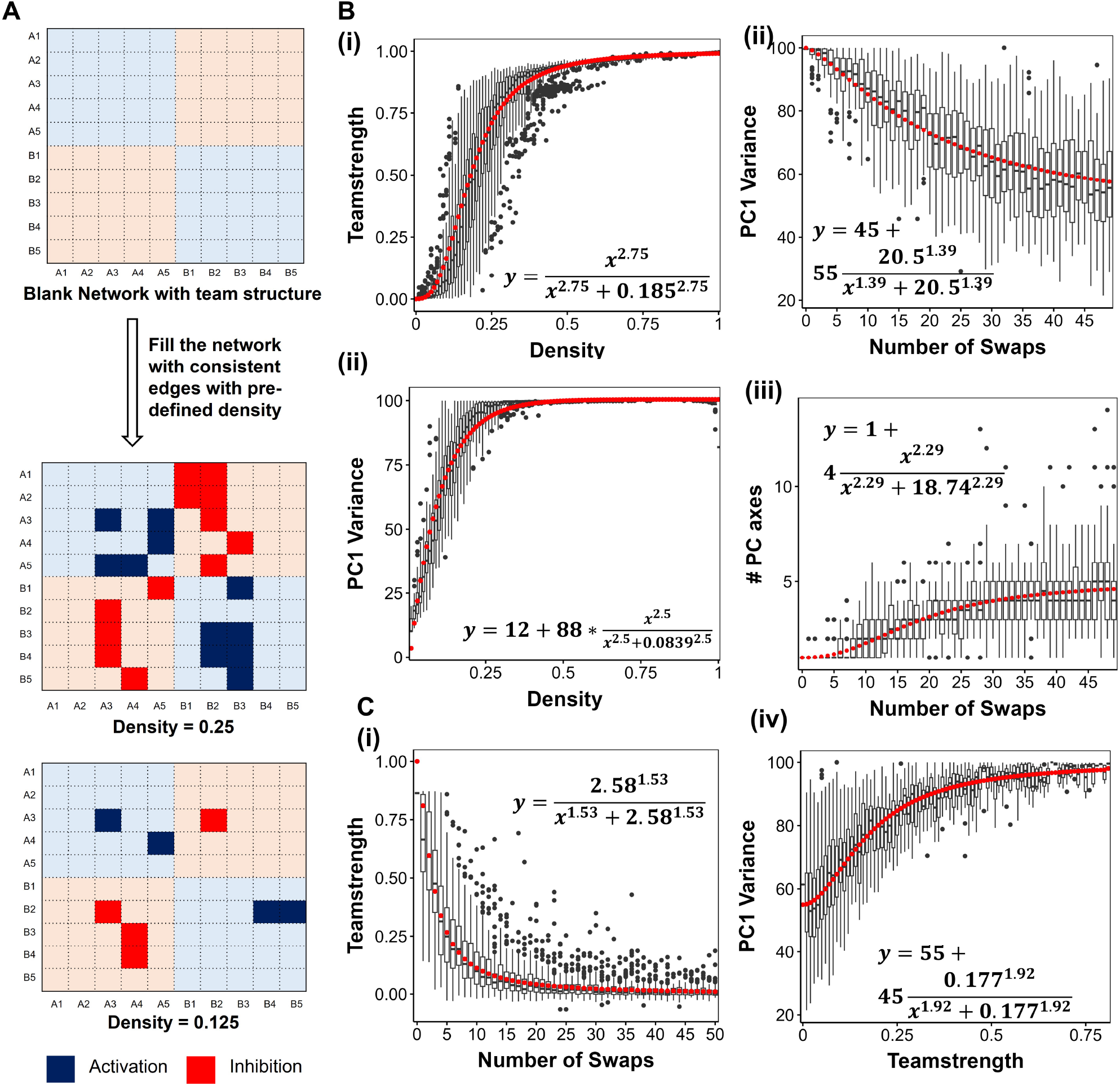
Artificial network analysis to validate the generalizability of the findings. **A**. Schematic for generating artificial networks with teams. **B**. Randomization of artificial networks. Box plots depicting the change in (i) Team strength, (ii) PC1 Variance, and (iii) Number of PC axes needed to explain 90% variance against the number of edges swapped in the artificial network. (iv) PC1 Variance against teamstrength of random networks. **C**. Analysis of artificial networks generated with different densities. Change in (i) PC1 Variance and (ii) Number of PC axes needed to explain 90% variance against network density. The fits are represented using red curves, and the corresponding equations have been reported in each case.

We started by studying the relationship between the density of the network and team strength, given the team structure. We find that the mean team strength increases sigmoidally with density, with a threshold of 0.185 (18.5%, Figure 6B, i). Note that most of the change happens only under a density limit of 0.3 (30%), which can serve as a cutoff on the density of binary cell-fate decision networks. Indeed, all the biological networks we have studied so far, including SCLC as being the most dense, obey this limit. The PC1 variance showed a sigmoidal increase with the density as well, with a similar Hill coefficient but hitting saturation earlier than that of team strength. On the other hand, the number of PCs saw a sigmoidal decrease with density (Figure S5).

We then applied randomization to the artificial networks. As seen with biological networks, we find the team strength of the networks decreasing with the number of swaps (Figure 6C i) and the dimensionality of the phenotypic space increasing, as seen by a decrease in the PC1 variance (Figure 6C ii) and an increase in the number of PC axes required to explain 90% variance (Figure 6C iii). Furthermore, we see that PC1 variance increases sigmoidally with teamstrength (Figure 6C iv) and the number of pc axes decrease with increasing teamstrength. Together, these results establish that the relationship between team strength and dimensionality of the phenotypic space is generalizable to a broad class of networks, further emphasizing teamstrength as a generalizable design principle that exerts strong control over the emergent phenotypic space.

## 3 Discussion

Cell-fate decisions are ubiquitous in biological systems. Cells dynamically change phenotypes in response to varying biochemical and biomechanical signals. During the development of an organism, starting from a totipotent stem cell, a sequence of decision-making events plays out, often through binary branching points, to facilitate cellular differentiation well-controlled in space and time [26]. The robustness of decision-making is frequently observed in developmental contexts, where phenotypes are sensitive only to a small set of specific perturbations, a property known as canalization [27]. Thus, it becomes imperative to understand how despite the size and complexity of regulatory networks underlying such phenotypic changes, only a few phenotypes are generated. Recent studies have begun to elucidate the design principles of underlying regulatory networks from various biological contexts: canalization [28], and giving rise to relatively simple, low-dimensional dynamics [16]. Our results suggest that this low dimensionality in phenotypic space is an emergent consequence of a latent structure in these regulatory networks - the presence of “teams” of nodes such that members within a team activate each other, and those across teams inhibit one another.

We had previously identified teams of nodes in regulatory networks driving the phenotypic decisions in EMP [5], SCLC [22] and melanoma [29]. We found that “teams” could lead to minimal frustration in biological networks. Recently, many biological networks were found to be minimally frustrated and capable of enabling a low-dimensional phenotypic space [16, 30]. Our results, therefore, provide a mechanistic understanding of these trends; we find that only in networks with high team strength most of the variance in the steady-state space (¿ 90%) is explained by PC1. By comparing the behavior of wild-type biological networks with their multiple randomized counterparts, we establish that the stronger the “teams”, the higher the variance explained by PC1. Thus, networks showing weak team strength have higher dimensional dynamics in phenotypic space, because it takes many more PCs together to be able to recapitulate the 90% variance.

This analysis is also strengthened by our calculations for two-team artificial networks, where we show how team strength and PC1 variance increase sigmoidally as a function of the density of edges in the network. By approximately 25% density, team strength and PC1 variance both show largely saturating behavior, suggesting that only one-fourth of edges connected in a two-team topology may be sufficient to allow for low dimensional dynamics in phenotypic space. Intriguingly, biological networks studied here show similar values of density in terms of the mean number of edges per node. Therefore, artificial networks provide an amenable tool to derive generalizable design principles. Future directions include identifying how impurities in network topology (i.e., members within a team inhibit each other and/or members across teams activate each other) or how the presence of more than two teams (e.g., as expected to be present in CD4+ T-cell differentiation [31]) changes these trends observed here. Overall, we establish that mutually inhibiting two “teams” of nodes is a design principle underlying the low dimensionality of phenotypic space seen experimentally.

Importantly, we detected this low dimensionality feature in corresponding transcriptomic data of SCLC and EMT-associated gene sets. This low dimensionality is maintained despite replacing many genes in the signatures with housekeeping genes, suggesting the presence of strong network topology features such as teams of nodes underlying the control of coordinated expression levels.

Our current analysis is limited to one axis of plasticity in each context, i.e., epithelial-mesenchymal, gonadal cell fate determination, and non-neuroendocrine/neuroendocrine cell fate determination. Cancer cells have been shown to exhibit multiple interconnected axes of phenotypic transitions (e.g., EMP, drug resistance, stemness, and metabolic plasticity [32, 33, 34, 35]. Studies suggest that these transitions can happen together (eg. EMT leading to the acquisition of stemness and/or drug resistance), suggesting interconnectivity between the underlying regulatory networks [36, 37, 38, 39]. An important direction for the current study is to understand how the low-dimensional dynamics scale across these interdependent axes of phenotypic heterogeneity and how the team structure evolves under such interactions.

## 4 Methods

### 4.1 RAndom CIrcuit PErturbaiton (RACIPE)

RACIPE [17] is a tool that simulates transcriptional regulatory networks (TRNs) in a continuous manner. Given a TRN, it constructs a system of Ordinary Differential Equations representing the network. For a given node *T* and a set of input nodes *P*_*i*_ and *N*_*j*_ that activate and inhibit *T* respectively, the corresponding differential equation takes the following form:

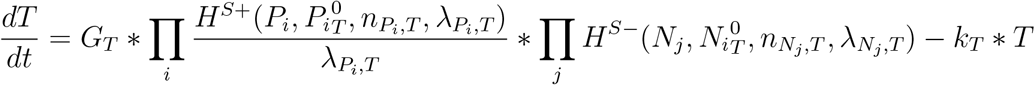

Here, *T, P*_*i*_ and *N*_*j*_ represent the concentrations of the species. *G*_*T*_ and *k*_*T*_ denote the production and degradation rates, respectively. 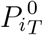 is the threshold value of *P*_*i*_ concentration at which the non-linearity in the dynamics of *T* due to *P*_*i*_ is seen. *n* is termed as Hill-coefficient and represents the extent of non-linearity in the regulation. *λ* represents the fold change in the target node concentration upon over-expression of regulating node. Finally, the functions *H*^*S*+^ and *H*^*S−*^ are known as shifted hill functions and represent the regulation of the target node by the regulatory node. The hill shift function takes the following form:

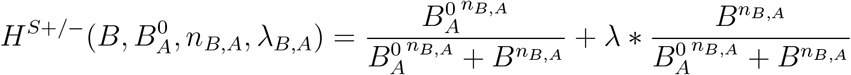

Note that, for high values of the regulatory node concentration, *H*^*S*+*/−*^ approaches *λ*.

For the model generated in this way, RACIPE randomly samples parameter sets from a pre-defined set of parameter ranges estimated from BioNumbers [40]. The ranges as reported by Huang et al [17] are as follows:

**Table 1.** Parameter ranges used by RACIPE.

| Parameters | Minimum | Maximum |
| --- | --- | --- |
| Production Rate ($G$) | 1 | 100 |
| Degradation Rate ($k$) | 0.1 | 1 |
| Fold Change (Inhibition $\lambda$) | 0.01 | 1 |
| Fold Change (Activation $\lambda$) | 1 | 100 |
| Hill Coefficient $n$ | 1 | 6 |
| Threshold | The ranges depend on inward regulations - half functional rule |  |

At each parameter set, RACIPE integrates the model from multiple initial conditions and obtains steady states in the initial condition space. The output, hence, comprises of the collection of parameter sets and corresponding steady states obtained from the model. For the current analysis, we used a sample size of 10000 for parameter sets and 100 for initial conditions. The parameters were sampled via a uniform distribution and the ODE integration was carried out using Euler’s method of numerical integration.

### 4.2 Processing RACIPE output

For a given network with *i* = [1, *n*] nodes, the steady state expression levels of the nodes were normalized in the following way:

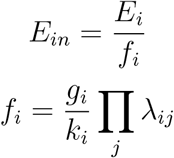

Where, for the *i*^*th*^ node, *E*_*in*_ is the normalized expression level, *E*_*i*_ is the steady state expression level, *f*_*i*_ is the normalization factor, *g*_*i*_ and *k*_*i*_ are production and degradation of the *i*^*th*^ node corresponding to the current steady state and *λ*_*ij*_ are the fold change in expression of *i* due to node *j* = [1, *n*]. The collection of these normalized steady states was used for PCA.

### 4.3 Boolean Simulations

We adapt the ising formalism of threshold-based boolean rules to simulate the artificial networks [14]. Briefly, each node is can have one of the two expression levels: *−*1 and +1 representing high or low expression, and each edge in the network (represented by *J*_*ik*_ for the edge from *i*^*th*^ to the *k*^*th*^ node) can have one of the three weights: *−*1, +1 and 0 for inhibition, activation and no interaction respectively. The update rule for the state of the *i*^*th*^ node at time *t, s*_*i*_(*t*) is as follows:

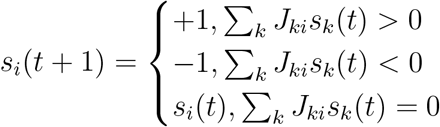

### 4.4 Network randomization

Network randomization is carried out by randomly swapping the edges in the network. Each iteration involves randomly selecting a pair of edges and swapping their edge type (activation/inhibition). For each random network, we carry out 10 to 100 such iterations. For each biological network, we generate 100 random networks.

### 4.5 Calculation of team strength

For each network, we first generate the adjacency matrix *A* = {*−*1, 0, 1}^*NXN*^, where *N* is the number of nodes in a network. Each cell of the adjacency matrix represents an edge, with 1 for inhibition, 1 for activation, and 0 for no interaction. We then generate the influence matrix using the following formula:

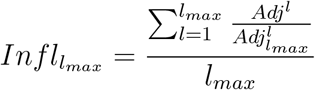

where *Adj*_*max*_ is derived by setting all non-zero entries of the adjacency matrix to 1 and the division is element-wise. We then take the sign of the influence matrix, augment the transpose of the influence matrix with itself, and apply hierarchical clustering to identify the two teams. We then calculate the team strength as follows:

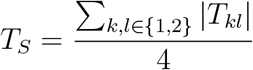

where

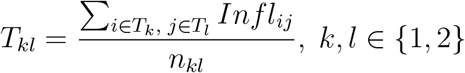

### 4.6 Disconnected nodes in biological networks

We generated networks with disconnected nodes by randomly selecting *n* nodes (0 ≤*n*≤*N*) and removing all incoming and outgoing edges connected to these nodes. Since RACIPE cannot simulate disconnected nodes, we added self-activation and self-inhibition to these nodes. For each *n* we generate 50 networks, representing 50 random choices of *n* nodes to be disconnected. We then simulate each network in RACIPE and estimate the PC1 variance and number PC axes.

### 4.7 Statistical analysis

Spearman correlation coefficients were calculated using the *cor*.*test* function from R 4.1. One-way ANOVA test was performed using the function *f_oneway* from the *scipy* package in Python 3.8.

## 5 Author Contributions

MKJ and KH designed research; KH, PH and TP analysed the data; AS, AG, PH and KH carried out simulations; all authors discussed results and participated in the preparation of the manuscript.

## 6 Conflict of Interest

The authors declare no competing Financial or non-Financial interests.

## 7 Data and Code Availability

The raw data generated for this study and derived datasets supporting the current findings are available from the corresponding author (MKJ, KH) upon reasonable request. The processed data files and the codes are available at the github repository: https://github.com/aashnasaxena/Teams-PC1.

## 8 Funding Information

KH is supported by the Prime Minister’s Research Fellowship (PMRF). MKJ was supported by Ramanujan Fellowship (SB/S2/RJN-049/2018) by the Science and Engineering Research Board (SERB), Department of Science and Technology, Government of India. HL was supported in part by the Center for Theoretical Biological Physics, NSF PHY-2019745.

## Supplementary Figures

**Figure S1:**
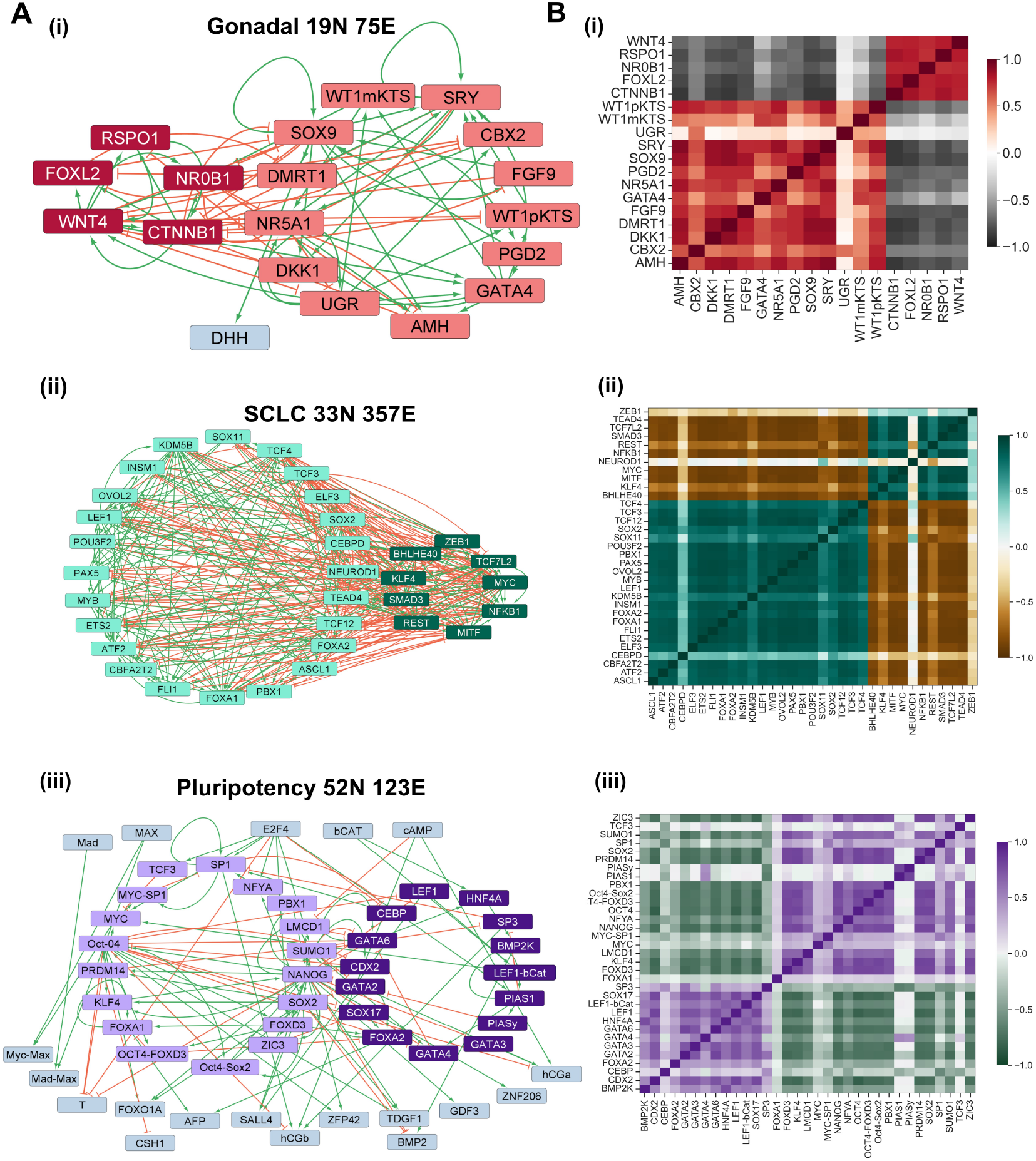
Cell fate decision networks. **A** Network diagrams for (i) Gonadal, (ii) SCLC, and (iii) iPSC network. **B** Correlation matrix depicting the pairwise correlations between the node expression levels across all parameter sets in RACIPE for (i) Gonadal, (ii) SCLC, and (iii) iPSC network.

**Figure S2:**
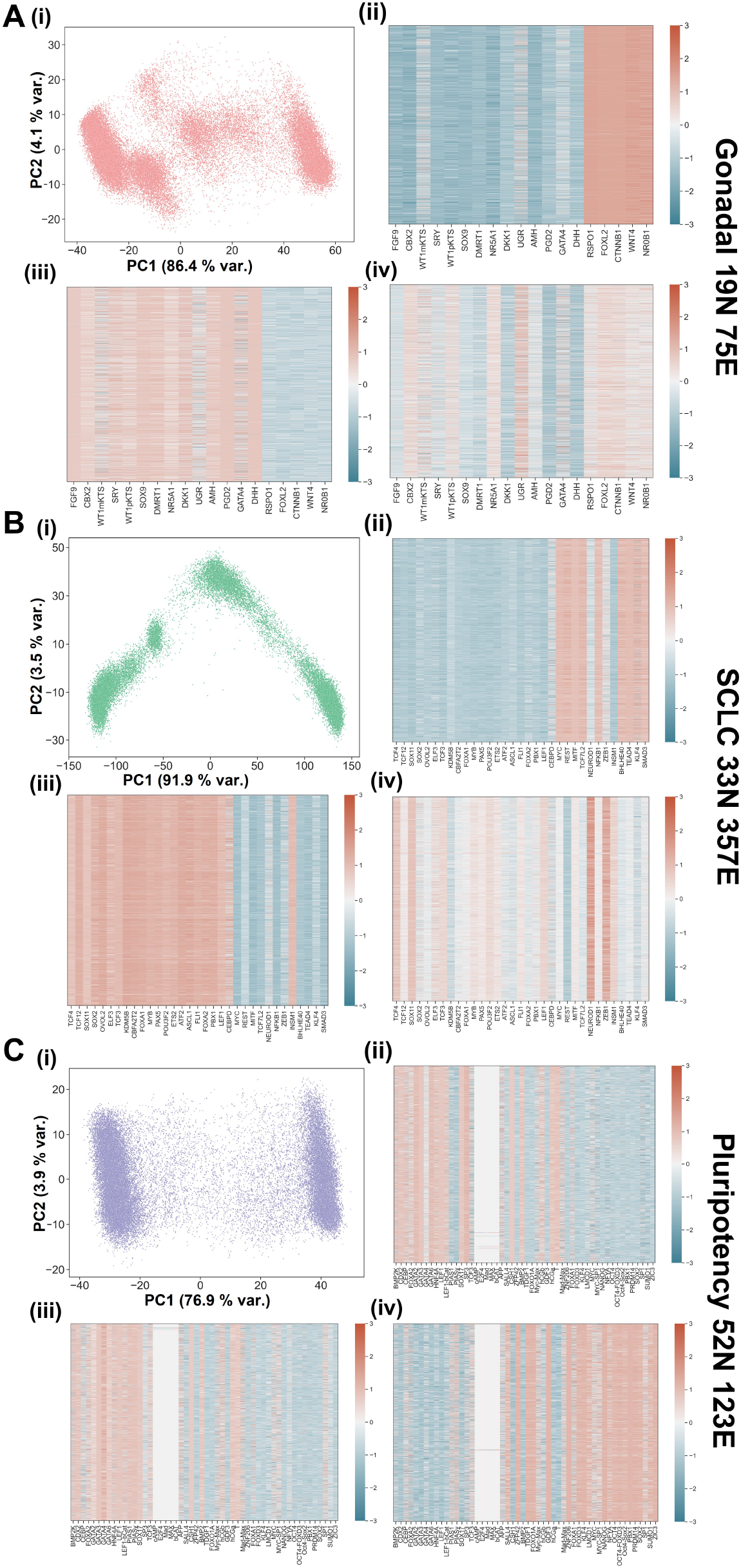
Structural similarities between teams and PC1. **A** Scatterplot mapping the solutions generated from RACIPE on the axes of PC1 and PC2. Heatmaps depicting the expression levels of the nodes of the network for each individual cluster Gonadal fate decision network. **B** Same as **A** but for SCLC network. **C** Same as **A** but for Pluripotency network.

**Figure S3:**
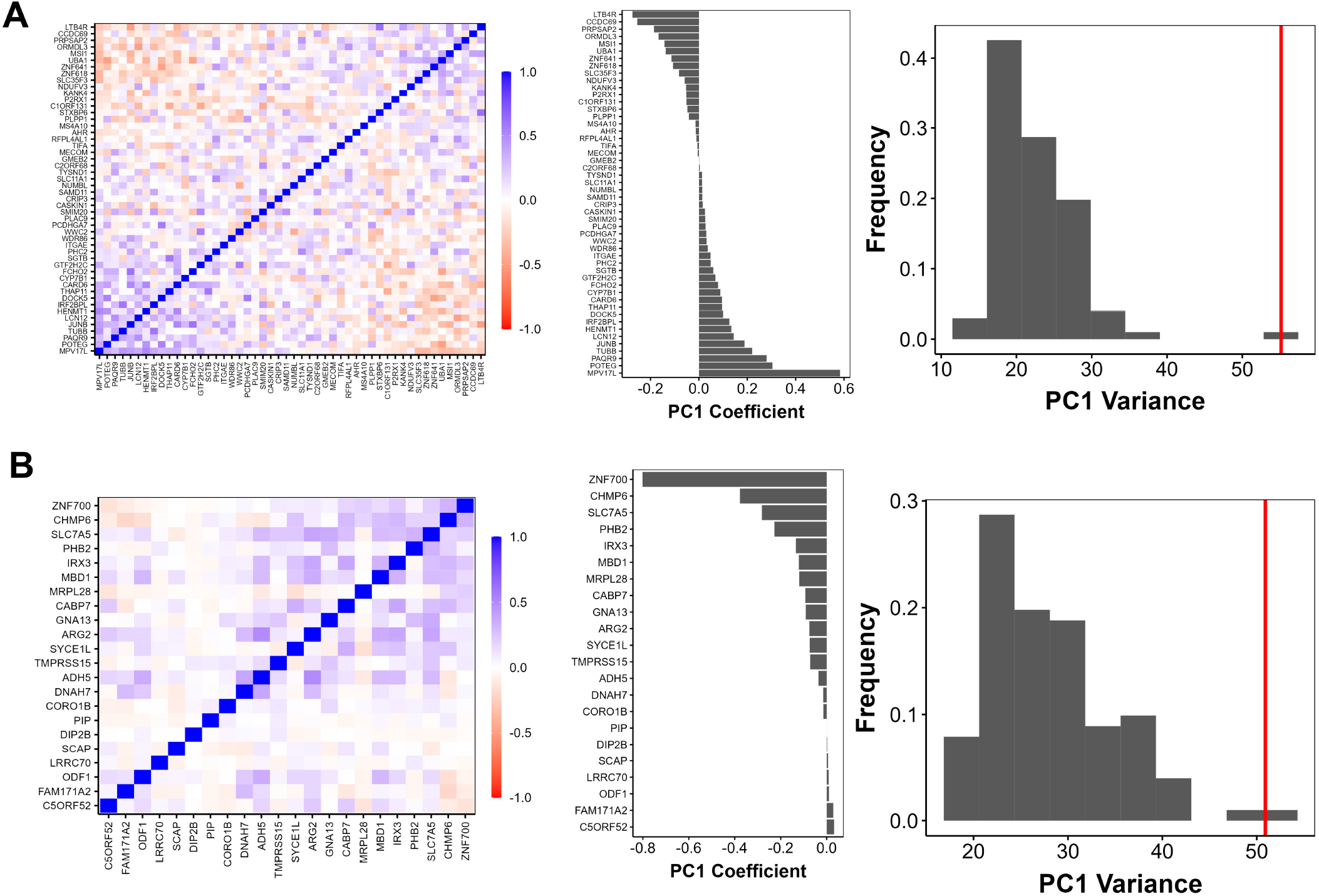
Dimensionality of transcriptional expression matrices of random sets of genes. **A**. (i) Pairwise correlation of expression for the 32 random genes in the CCLE SCLC cell-line data. (ii). Coefficients of first principle component axis for the geneset-cell line combination in (i). (iii) Distribution of PC1 variance for 100 choices of 32 random genes. PC1 Variance corresponding to SCLC gene set is represented by red vertical line. **B**. Same as A, but for CCLE non-SCLC cell-line data.

**Figure S4:**
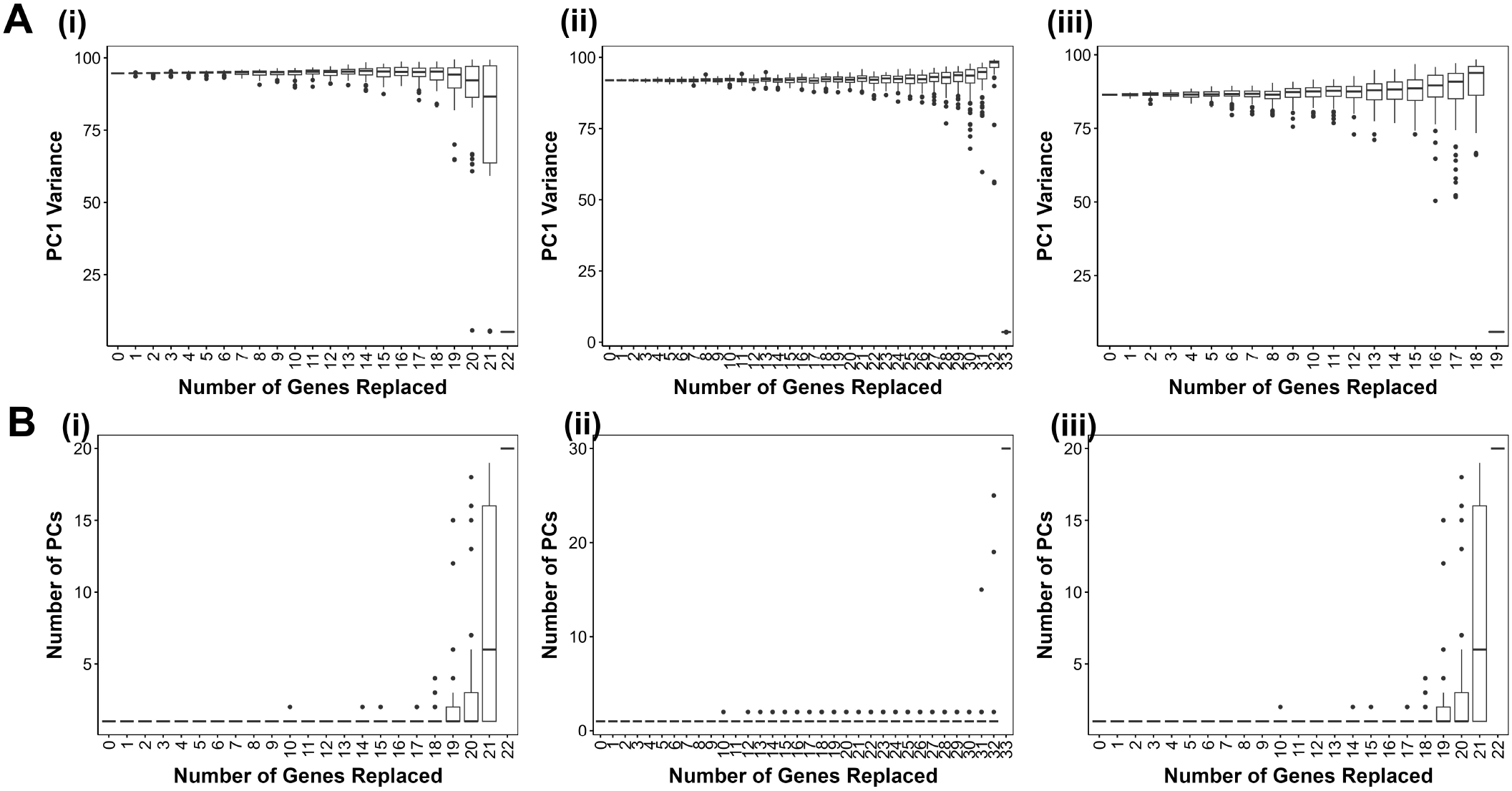
Node replacement analysis on biological networks. **A**. PC1 variance against the number of nodes replaced for (i) EMT, (ii) SCLC and (iii) gonadal networks. **B**. Same as A, but for number of PC axes.

**Figure S5:**
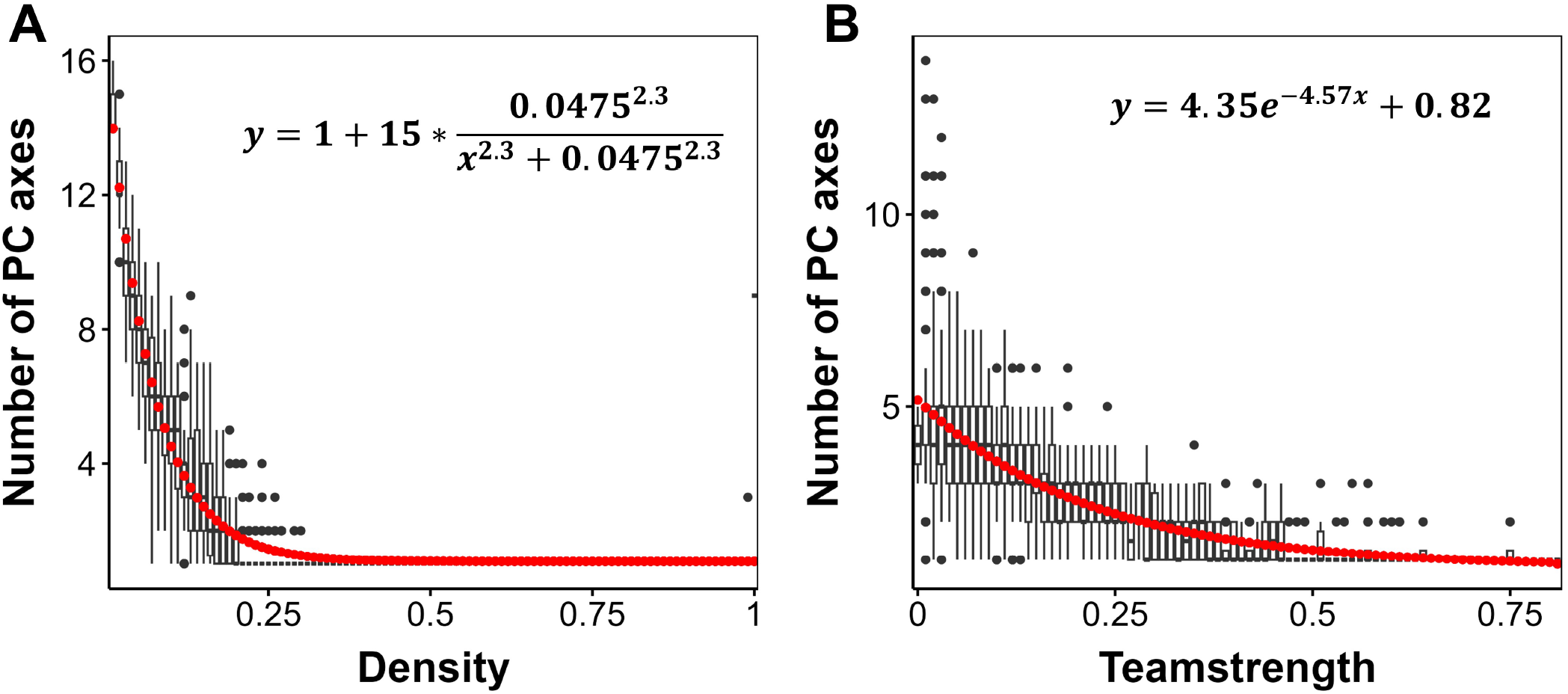
Analysis of Artificial networks. Number of PC axes against **A**. Density and **B**. Teamstrength for artificial networks.

## Notes

### Competing Interest Statement

The authors have declared no competing interest.

### Summary of Updates

New results added - Figure 4-6 and corresponding main text and SI figures

## References

[1] Vickaryous MK, Hall BK (2006) Human cell type diversity, evolution, development, and classification with special reference to cells derived from the neural crest. Biological Reviews 81(03):425.

[2] Jones RC, et al. (2022) The tabula sapiens: A multiple-organ, single-cell transcriptomic atlas of humans. Science 376(6594):eabl4896.

[3] Gulati GS, et al. (2020) Single-cell transcriptional diversity is a hallmark of developmental potential. Science 367(6476):405–411.

[4] Jolly MK, Ware KE, Gilja S, Somarelli JA, Levine H (2017) Emt and met: necessary or permissive for metastasis? Molecular Oncology 11(7):755–769.

[5] Hari K, Ullanat V, Balasubramanian A, Gopalan A, Jolly MK (2022) Landscape of epithelial–mesenchymal plasticity as an emergent property of coordinated teams in regulatory networks. eLife 11:e76535.

[6] Hong T, et al. (2015) An ovol2-zeb1 mutual inhibitory circuit governs bidirectional and multistep transition between epithelial and mesenchymal states. PLOS Computational Biology 11(11):e1004569.

[7] Roca H, et al. (2013) Transcription factors OVOL1 and OVOL2 induce the mesenchymal to epithelial transition in human cancer. PLoS ONE 8(10):e76773.

[8] Zhou JX, Huang S (2011) Understanding gene circuits at cell-fate branch points for rational cell reprogramming. Trends in Genetics 27(2):55–62.

[9] Pastushenko I, et al. (2018) Identification of the tumour transition states occurring during EMT. Nature 556(7702):463–468.

[10] Jolly MK, et al. (2019) Hybrid epithelial/mesenchymal phenotypes promote metastasis and therapy resistance across carcinomas. Pharmacology & Therapeutics 194:161–184.

[11] Lu M, Jolly MK, Levine H, Onuchic JN, Ben-Jacob E (2013) MicroRNA-based regulation of epithelial–hybrid–mesenchymal fate determination. Proceedings of the National Academy of Sciences 110(45):18144–18149.

[12] Tian XJ, Zhang H, Xing J (2013) Coupled reversible and irreversible bistable switches underlying TGF-induced epithelial to mesenchymal transition. Biophysical Journal 105(4):1079–1089.

[13] Subbalakshmi AR, Ashraf B, Jolly MK (2022) Biophysical and biochemical attributes of hybrid epithelial/mesenchymal phenotypes. Physical Biology 19(2):025001.

[14] Font-Clos F, Zapperi S, Porta CAML (2018) Topography of epithelial–mesenchymal plasticity. Proceedings of the National Academy of Sciences 115(23):5902–5907.

[15] Steinway SN, et al. (2015) Combinatorial interventions inhibit TGFβ-driven epithelial-to-mesenchymal transition and support hybrid cellular phenotypes. npj Systems Biology and Applications 1:15014.

[16] Tripathi S, Kessler DA, Levine H (2023) Minimal frustration underlies the usefulness of incomplete regulatory network models in biology. Proceedings of the National Academy of Sciences 120(1):e2216109120.

[17] Huang B, et al. (2017) Interrogating the topological robustness of gene regulatory circuits by randomization. PLOS Computational Biology 13(3):e1005456.

[18] Udyavar AR, et al. (2017) Novel Hybrid Phenotype Revealed in Small Cell Lung Cancer by a Transcription Factor Network Model That Can Explain Tumor Heterogeneity. Cancer Res 77(5):1063–1074.

[19] Chang R, Shoemaker R, Wang W (2011) Systematic search for recipes to generate induced pluripotent stem cells. PLoS Comput Biol 7(12):e1002300.

[20] Ríos O, et al. (2015) A boolean network model of human gonadal sex determination. Theor Biol Med Model 12(1):26.

[21] Barretina J, et al. (2012) The cancer cell line encyclopedia enables predictive modelling of anticancer drug sensitivity. Nature 483(7391):603–607.

[22] Chauhan L, Ram U, Hari K, Jolly MK (2021) Topological signatures in regulatory network enable phenotypic heterogeneity in small cell lung cancer. eLife 10:e64522.

[23] Lissa D, et al. (2022) Heterogeneity of neuroendocrine transcriptional states in metastatic small cell lung cancers and patient-derived models. Nature Communications 13(1).

[24] Aiello NM, et al. (2018) EMT subtype influences epithelial plasticity and mode of cell migration. Developmental Cell 45(6):681–695.e4.

[25] Eisenberg E, Levanon EY (2013) Human housekeeping genes, revisited. Trends in Genetics 29(10):569–574.

[26] Agozzino L, Balazsi G, Wang J, Dill KA (2020) How do cells adapt? stories told in landscapes. Annual Review of Chemical and Biomolecular Engineering 11(1):155–182.

[27] Du Z, et al. (2015) The regulatory landscape of lineage differentiation in a metazoan embryo. Developmental Cell 34(5):592–607.

[28] Kadelka C, Butrie TM, Hilton E, Kinseth J, Serdarevic H (2020) A meta-analysis of boolean network models reveals design principles of gene regulatory networks. arXiv:2009.01216.

[29] Pillai M, Jolly MK (2021) Systems-level network modeling deciphers the master regulators of phenotypic plasticity and heterogeneity in melanoma. iScience 24(10):103111.

[30] Tripathi S, Kessler DA, Levine H (2020) Biological networks regulating cell fate choice are minimally frustrated. Physical Review Letters 125(8).

[31] Duddu AS, Sahoo S, Hati S, Jhunjhunwala S, Jolly MK (2020) Multi-stability in cellular differentiation enabled by a network of three mutually repressing master regulators. Journal of The Royal Society Interface 17(170):20200631.

[32] Thankamony AP, Saxena K, Murali R, Jolly MK, Nair R (2020) Cancer stem cell plasticity – a deadly deal. Frontiers in Molecular Biosciences 7:79.

[33] Jia D, et al. (2019) Elucidating cancer metabolic plasticity by coupling gene regulation with metabolic pathways. Proceedings of the National Academy of Sciences 116(9):3909–3918.

[34] Vipparthi K, et al. (2022) Emergence of hybrid states of stem-like cancer cells correlates with poor prognosis in oral cancer. iScience 25(5):104317.

[35] Bhatia, et al. (2019) Interrogation of phenotypic plasticity between epithelial and mesenchymal states in breast cancer. Journal of Clinical Medicine 8(6):893.

[36] Sahoo S, et al. (2021) A mechanistic model captures the emergence and implications of non-genetic heterogeneity and reversible drug resistance in ER breast cancer cells. NAR Cancer 3(3):zcab027.

[37] Galbraith M, Levine H, Onuchic JN, Jia D (2023) Decoding the coupled decision-making of the epithelial-mesenchymal transition and metabolic reprogramming in cancer. iScience 26(1):105719.

[38] Pasani S, Sahoo S, Jolly MK (2020) Hybrid e/m phenotype(s) and stemness: A mechanistic connection embedded in network topology. Journal of Clinical Medicine 10(1):60.

[39] Biddle A, Gammon L, Liang X, Costea DE, Mackenzie IC (2016) Phenotypic plasticity determines cancer stem cell therapeutic resistance in oral squamous cell carcinoma. EBioMedicine 4:138–145.

[40] Milo R, Jorgensen P, Moran U, Weber G, Springer M (2009) BioNumbers—the database of key numbers in molecular and cell biology. Nucleic Acids Research 38(suppl 1):D750–D753.

